# Epigenetics Insights from Perceived Facial Aging

**DOI:** 10.1101/2023.05.29.542727

**Authors:** Klemo Vladimir, Marija Majda Perišić, Mario Štorga, Ali Mostashari, Raya Khanin

## Abstract

Facial aging is the most visible manifestation of aging. People desire to look younger than others of the same chronological age. Hence, perceived age is often used as a visible marker of aging, while biological age, often estimated by methylation markers, is used as an objective measure of age. Multiple epigenetics-based clocks have been developed for accurate estimation of general biological age and the age of specific organs, including the skin. However, it is not clear whether the epigenetic biomarkers (CpGs) used in these clocks are drivers of aging processes or consequences of aging.

In this proof-of-concept study, we integrate data from GWAS on perceived facial aging, and EWAS on CpGs measured in blood. By running EW Mendelian randomization, we identify hundreds of putative CpGs that are potentially causal to perceived facial aging with similar numbers of damaging markers that causally drive or accelerate facial aging and protective methylation markers that causally slow down or protect from aging. We further demonstrate that while candidate causal CpGs have little overlap with known epigenetics-based clocks, they affect genes or proteins with known functions in skin aging such as skin pigmentation, elastin, and collagen levels. Overall, our results suggest that blood methylation markers reflect facial aging processes, and thus can be used to quantify skin aging and develop anti-aging solutions that target the root causes of aging.

## 1 Introduction

Aging is a multifactorial process. People age at different rates, and chronological age does not always reflect the actual biological age of a person. Therefore, the evaluation of biological age is of high relevance for understanding aging processes and developing interventions to extend healthy aging. Numerous biological clocks based on high-throughput biomarkers such as methylation markers, gene expression, protein levels, and microbiota, have been proposed to measure a person’s biological age.

The ‘first generation’ of biological clocks have been trained on chronological age, while the ‘second generation’ of clocks utilize other age-related measures, including time to all-cause mortality or biomarkers of morbidity. Many biological clocks demonstrate high accuracy in predicting a person’s chronological age or age-related measures [1–6]. Novel computational solutions bolster the reliability of biological clocks by extracting the shared aging signal across many CpGs while minimizing noise from individual CpGs [7].

Epigenetic predictors have become the gold standard biomarker for the estimation of human biological age. They are highly reproducible and have been validated in multiple studies. Most existing epigenetics-based biological clocks are developed using penalized regression approaches such as elastic net on a training dataset yielding a subset of CpG (from a few dozen to a few hundred) that are organized into a weighted linear prediction model of biological age. The discrepancy between epigenetic-based biological age and chronological age called age acceleration is predictive of all-cause mortality and various age-associated diseases.

Skin samples have been utilized to construct various biological clocks. Firstly, the multi-tissue age predictor based on 353 CpGs was developed using 8,000 samples encompassing 51 healthy tissues and cell types including skin [4]. This pan-tissue predictor can be used to estimate the age of most human cell types, tissues, and organs. The skin-blood biological clock utilizes 391 CpGs and it outperforms the pan-tissue clock in all metrics of accuracy in skin samples (including fibroblasts, keratinocytes, and endothelial cells) and blood samples [6]. More recently, a skin-specific biological clock was developed using data from 508 human skin samples, including cultured skin cells and human skin biopsies [8]. This clock was further utilized for identifying a senolytic compound that reduces biological age [9].

However, it is not clear whether CpG sites used in these biological clocks are just convenient markers, consequences of aging, or active drivers of aging [10, 11]. Current biological clocks utilize epigenetic biomarkers that are merely correlated with chronological age, and the model that optimally predicts an outcome from a statistical perspective is selected. Generally, no preference is given to the location or possible biological role of the CpG-features used as inputs in the machine learning procedure. While some biological clocks demonstrate high accuracy in predicting a person’s chronological age or age-related measures, the molecular mechanisms of how these clocks work are not clear. There is little overlap between CpGs in different biological clocks pointing to the significant redundancy in DNA methylation.

It is therefore critical to identify causal epigenetic biomarkers that are not simply correlated with chronological age, or age-related measures but also reflect either damage that accumulates with age or anti-aging protection. This would enable more efficient development of preventative anti-aging measures that slow down, or even reverse, age-related damage while inducing protective anti-aging processes.

In this proof-of-concept study, we explore a hypothesis that CpGs causally associated with perceived facial aging reflect biological processes that underlie skin aging. Perceived facial age is a highly relevant marker of aging. It has been shown to be associated with all-cause mortality, independent of chronological age, and other aging-associated traits [12–15]. Usually, perceived facial age is determined by specialists such as dermatologists, or by other people. Several studies reported that large-scale data on perceived facial age can provide insights into biological age-related processes. For example, a genome-wide association study of adult participants of the UK Biobank identified 74 independently associated genetic loci which were enriched for cell signaling pathways, including the NEK6 and SMAD2 subnetworks [16]. The NEK6 subnetwork helps govern the initiation of mitosis and progression through the cell cycle and prevents cell senescence. Functional analysis of genes associated with self-perceived facial age revealed a significant over-representation of skin pigmentation, extracellular matrix, and immune-related pathways that are correlated with facial aging [17].

Recalling that “correlation is not causation “, we set out to identify CpG sites that exert a causal effect on facial aging. To this end, we integrated genome-wide association (GWAS) data on perceived facial aging of UKBB participants with epigenome-wide methylation quantitative loci (meQTL) from blood, and run Epigenome-Wide Mendelian Randomization (EWMR). We found that methylation markers that causally affect facial aging have little or no overlap with most existing epigenetics-based biological clocks. However, causal CpG sites provide valuable insights into molecular mechanisms of skin aging. Furthermore, our results suggest that blood methylation markers reflect aging processes in the skin, and hence can be utilized to quantify skin aging and to potentially evaluate and develop anti-aging skin treatments.

## 2 Methods

### 2.1 Mendelian Randomization

#### 2.1.1 EWMR

As exposures in the Mendelian Randomization (MR), we utilized 11, 165, 559 SNP-CpG associations (methylation quantitative trait loci, meQTL) from whole blood samples identified through genome-wide association analysis from 3, 799 Europeans and 3, 195 South Asians [18]. In this study, we used results from the European cohort using genetic instruments at p-value threshold = 10^−6^.

Epigenome-wide MR (EWMR) was run on each CpG (exposure) - GWAS (outcome) pair. For the outcomes, facial aging (ukb-b-2148), telomere length (ieu-b-4879) were obtained from the `MR-Base` GWAS catalog [19]. Additional skin-related outcomes include actinic keratosis (finn-b-L12_ACTINKERA), acute skin changes due to ultraviolet radiation (finn-b-L12_OTHACUSKINSU, finn-b-L12 OTHACUSKINSUNNAS), facial pigmentation (ebi-a-GCST90002283), basal cell carcinoma (ebi-a-GCST90013410). For facial aging and telomere length, EWMR anayses were run on all available CpGs [18]. For other skin-related traits, EWMR analyses were run on CpGs identified as causal (*p <* 0.001) in the EWMR for facial aging.

To run EWMR we used the `TwoSampleMR` package in `R` statistical language. Pleiotropy was evaluated based on the intercept calculated by MR-Egger regression using mr_pleiotropy_test with p-value threshold *p* = 0.05. Exposure-outcome relationships that change by at least 5% in the odds ratio (*OR* ≥ 1.05 or *OR* ≤0.95) are reported.

#### 2.1.2 Other MR

EW-GE MR analysis evaluates relationships between CpGs (exposure) and gene expression (outcome). For each CpG and its nearby gene pair, EW-GE MR evaluates whether a CpG causally results in either an increase or decrease of expression of the gene. For this study, EW-GE MR was run for CpGs causal to facial aging as identified in the EWMR step. As above, methylation at a CpG site is proxied by meQTL genetic instruments and gene expression (GE) is proxied by eQTLs from the GTeX data for all tissues as available in the `TwoSampleMR` R package.

To further evaluate whether changes in gene expression (GE) causally affect facial aging (outcome), we ran for genes with available eQTLs for skin tissues (skin not sun-exposed and skin sun-exposed) and whole blood as available in the `TwoSampleMR` R package (GEMR).

### 2.2 Functional enrichment analysis

Enrichment analysis of causal CpGs was run using the `methylglm` function from the `methylgsa R` package. The `methylglm` function implements logistic regression adjusting for the number of probes in the enrichment analysis. It was run with default parameters using gene ontologies (GO, KEGG, and Reactome). Enrichment analysis on gene and protein levels was run using the FUMA Functional Mapping (https://fuma.ctglab.nl) using the default settings for all coding and noncoding genes. Gene-sets identified at FDR= 0.05 are reported as over-represented.

To assess relationships between CpG and serum protein levels, we used the EWAS catalog (http://www.ewascatalog.org).

## 3 Results

### 3.1 CpGs causally affect facial aging

Mendelian randomization (MR) is an established genetic computational approach for causal inference that recapitulates the principle of a randomized clinical trial (RCT). RCTs are very time-consuming and costly as they generally require a direct comparison of the cases and the controls to evaluate the effect of treatment (exposure). The idea behind the MR methodology is that genetic variants (SNPs), randomly assigned at conception, are robustly associated with the exposure and outcome, and not biased by environmental confounders. Thus, the biases can be circumvented by leveraging SNPs as instrumental variables (genetic instruments) to detect causal effects and estimate their magnitude.

The Epigenome-Wide Mendelian Randomization (EWMR) has been used to identify methylation markers that exert causal effects on several complex traits [20, 21], and general aging [11]. The methylation level at each CpG (exposure) is proxied by methylation quantitative trait loci (meQTL) as genetic instruments for exposure. In this study, we ran EWMR utilizing publicly available summary statistics data on perceived facial aging, telomere length, and other skin-related phenotypic traits as an outcome and meQTLs measured in the whole blood as an exposure (Figure 1 and Methods).

**Fig. 1.**
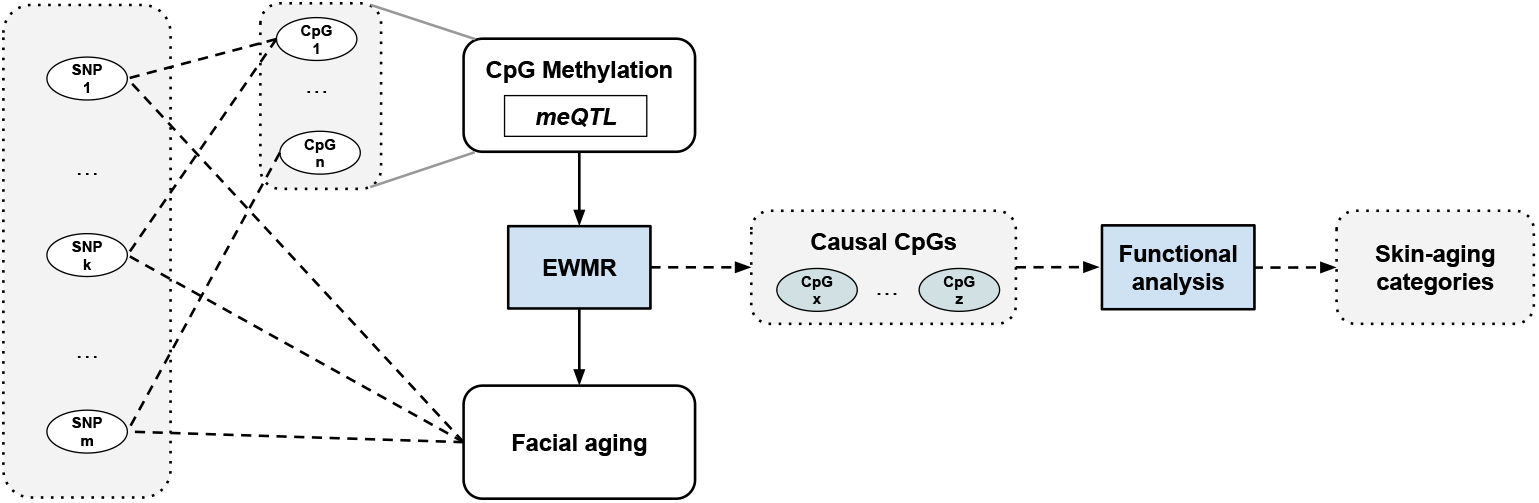
High-level illustration of the study design. Epigenome-Wide Mendelian Randomization (EWMR) is run to identify causal CpG sites. Epigenome-wide methylation quantitative loci (meQTL) from blood are used as exposure, and summary statistics from genome-wide association study (GWAS) on perceived facial aging is used as an outcome. EWMR is also run on telomere length, and several skin-related traits (see Methods). Functional analysis of CpGs causal to facial aging provides valuable insights into skin aging mechanisms.

Further, integration of eQTL data suggested that genetic variants, responsible for highly significant meQTL associations, also influence gene expression, pointing to a coordinated system of effects that are consistent with causality [20]. Large-scale studies reported that methylation at gene promoters is negatively correlated with gene expression, while methylation within gene bodies can be positively or negatively correlated with gene expression [22]. Furthermore, CpG sites typically regulate multiple transcripts, and while found to predominantly decrease gene expression, this was only the case for 53.4% across ≈47,000 significant CpG-transcript pairs [21]. These authors further demonstrated that GTeX data could be utilized as an outcome in the MR procedure wherein expression of a gene proxied by eQTLs across tissues, or in a specific tissue (e.g. skin), while methylation at a CpG site, proxied by genetic instruments, is used as an exposure. Thus, EW-GE MR assesses the causal effect of methylation at a specific CpG site on gene expression. Similarly, to identify the causal effect of gene expression (exposure) on a phenotypic trait, such as facial aging, the GEMR procedure is used.

Our EWMR analysis yielded 1299 candidate CpG sites (*p <* 0.001) that causally affect perceived facial aging (Methods) by at least 5% (OR*>*= 1.05 or OR*<*= 0.95) (Table S1). There are approximately similar numbers of damaging (659) and protective (640) CpGs (Figure 2). Damaging, or age-accelerating, CpGs causally increase or accelerate facial aging while protective, or adaptive, CpGs are causally associated with slower facial aging. Protective CpGs may also be referred to as promoting longevity or youthfulness methylation markers. This finding is in line with a recent study on general aging [11] suggesting that biological processes that adapt with age, or protect from early aging, at the epigenetic level are nearly as common as those mechanisms that drive or accelerate aging.

**Fig. 2.**
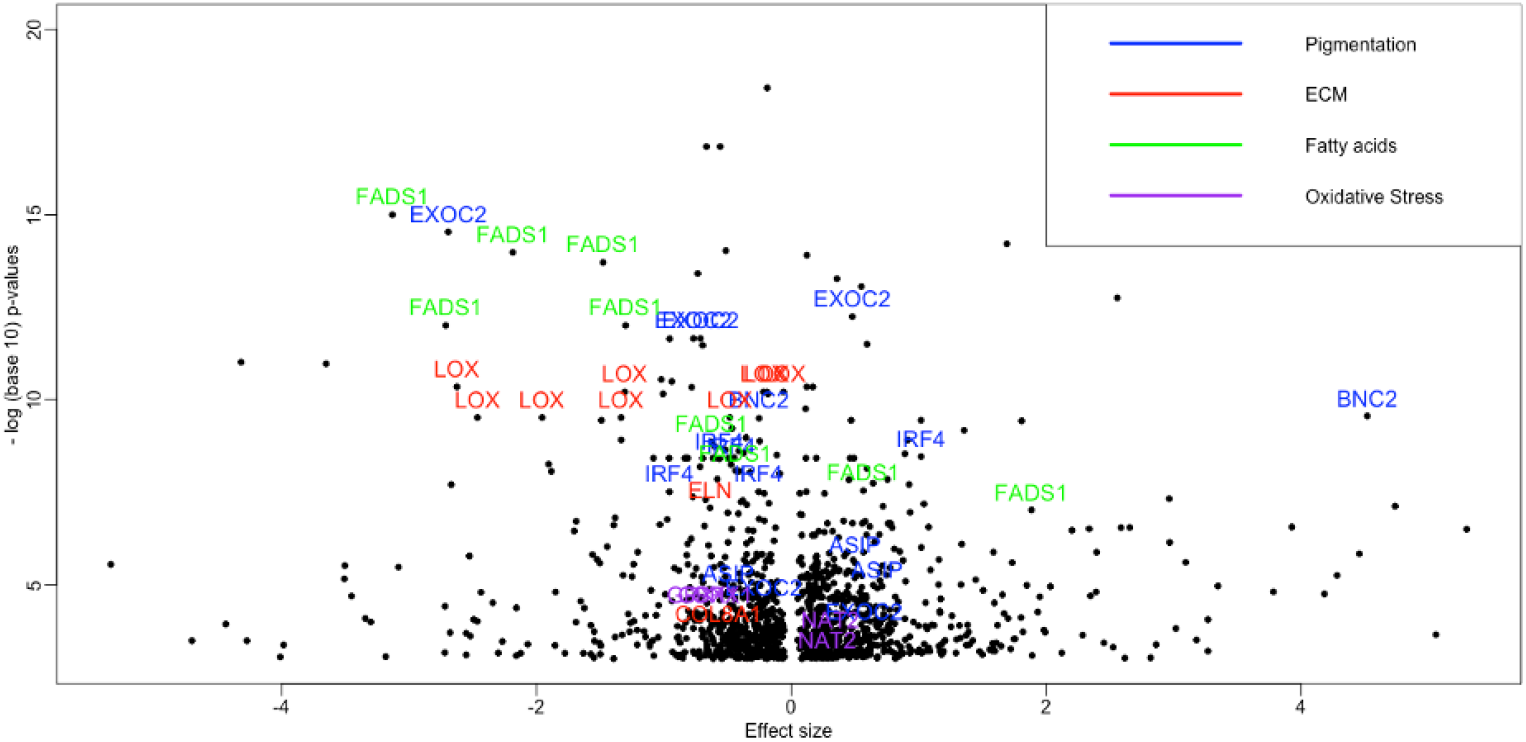
Volcano plot with effect sizes and p-values (*p <* 0.001) from EWMR analyses. Several representative genes from categories related to skin aging are shown. Pigmentation category includes genes IRF4, BNC2, EXOC2, ASIP/RALY. The extracellular matrix category includes genes COL8A1, ELN, LOX; the fatty acids group contains FADS1 and FADS2; the oxidative stress group includes NAT2 and GPX1.

Acknowledging that using unadjusted p-values (*p* ≤0.001) may yield false positive candidate CpGs, we perform various analyses to investigate whether blood-measured CpGs confer information on facial aging mechanisms (Figure 2). To this end, we first look at overlaps between causal CpGs identified in the EWMR analysis and the published biological clocks. We further perform functional enrichment analyses on the level of causal CpGs, and genes that annotate these CpGs. We also identify proteins reported to be associated with these CpGs in EWAS studies and followed up with functional analyses on the protein level.

### 3.2 Little overlap with epigenetic biological clocks

A number of published epigenetics-based clocks are highly predictive of biological age. Interestingly, only a small subset of CpGs are common among biological clocks, while CpG sites that approximate telomere length (DNAmTL) are entirely unique. This may indicate significant redundancy in DNA methylation.

To investigate the overlaps between candidate CpGs causal to facial aging and existing biological clocks we used the `methylCIPHER` package in `R` [23]. Overall, there is little, or no, overlap between CpGs from epigenetic-based biological clocks and CpGs potentially causal to facial aging. The largest overlap of CpGs causal to facial aging (8 CpGs) is with the updated PhenoAge clock (HRSInChPhenoAge). The effects of these CpGs in the updated PhenoAge clock and their causal effects in our study are anti-correlated (r= − 0.44). The second largest overlap (7 CpGs) is with the epigenetic-based predictor of chronological age [24] trained on 13, 661 methylation samples from blood and saliva [24].

Only 4 CpGs (cg03473532, cg10586358, cg10729426, cg06458239) from the Horvarth skin and blood age predictor [6] are identified as causal to facial aging in our EWMR analysis. Other biological clocks from the `methylCIPHER` package have 2 or 1 CpG in common with the list of causal CpG, while there are no overlaps with the majority of existing biological clocks. This suggests that existing epigenetics-based clocks built using correlated methylation markers are unlikely to reflect early causal age-driving or age-protective events. Similarly, out of the top 1000 CpGs reported to be correlated with facial aging of participants of the Lothian Birth Cohort, 1921 [25], only three CpGs are also identified as causal to facial aging.

The lack of overlap between CpGs causal to aging and epigenetics-based clocks is in line with the recent work on general aging [11]. These authors further suggested that although some epigenetics-based clocks contain CpGs causal to aging, they, by design, favor CpG sites with a higher correlation with age, and thus are not enriched with causal CpGs.

### 3.3 Annotation and Functional analysis of causal CpGs

Earlier studies reported that the majority of cross-tissue meQTLs showed consistent signs of cis-acting effects across tissues [2, 26] supporting the idea that genetically controlled CpGs are more likely to have common function across tissues compared to those less under direct genetic control. At the same time, a large portion of meQTLs shows tissue-specific characteristics, especially in the brain. To investigate molecular mechanisms underlying facial aging, we performed functional analyses on the level of CpGs inferred from the blood data, as well as on the level of genes and proteins.

Firstly, functional analysis of 1299 candidate CpGs causal to facial aging reveals several highly significantly over-represented GO categories that are known to play critical roles in skin aging. These include semaphorins, DNA repair, elastic fiber, and collagen related gene-sets, mitochondria membrane gene-sets, metabolic processes, and transmembrane transports of vitamins (e.g. thiamine) [Figure 3, Table S2].

**Fig. 3.**
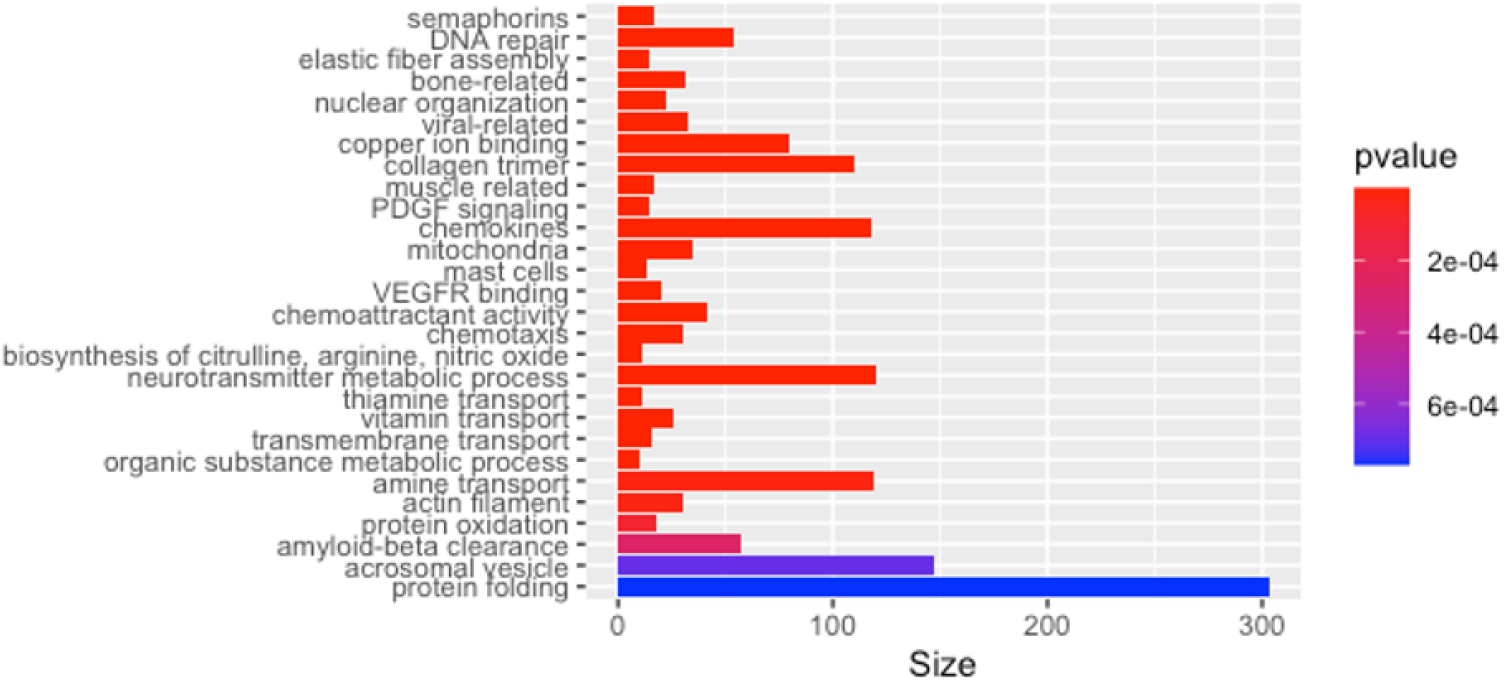
Bar-plot of adjusted p-values from the functional analysis of CpGs causal to facial aging. Full data are in Table S2. Similar categories are lumped together. For example, all significant semaphorin-related gene-sets are shown as one with the most significant adjusted p-value. The analysis is performed using `methylglm` function from the `methylgsa R` package.

Considered separately, damaging CpGs are enriched in collagen binding, multiple mitochondria-related gene-sets, vitamin (thiamine) transmembrane transport, hair cell differentiation, and several others [Table S2]. Protective CpGs are enriched in collagen formation, elastic fiber assembly, DNA biosynthetic process, Platelet-derived growth factor receptor (PDGFR) signaling, bone, muscle, and neuron development processes. Also, response to chemokines is significantly over-represented in protective CpGs. Chemokines are signaling proteins with roles in morphogenesis and wound healing. From an immunological perspective, chemokines and cytokines act in intercellular communicating immunity in skin aging [27].

To get further insight into the mechanistic functions, we first annotated all causal CpGs with nearby genes [Table S1]. This resulted in 412 (unique) genes that correspond to damaging CpGs, and 400 genes that correspond to protective CpGs, with a large overlap (132 genes) between the two lists. The reason for this overlap is that a methylation marker may increase the expression of a nearby gene, decrease it, or have no effect at all, depending on the genomic context [21]. Removal of overlapping genes yields 280 genes that correspond to damaging CpGs, and 268 genes that annotate protective CpGs. We further conducted an EW-GE MR study and identified several plausible mechanisms on how methylation at specific CpG site causally affect gene expression which in turn affect facial aging. In general, more high-throughput studies and additional analyses are needed to infer directions and sizes of the effect of methylation at CpG sites on gene expression.

Additionally, we utilized the EWAS catalog to connect causal CpGs with proteins whose levels are significantly associated with CpG sites [Table S1]. There is no available data to deduce whether associations between CpGs and protein levels are causal, or whether changes in protein levels drive aging or are the consequences of aging. However, by integrating the results of EWMR analysis and data from the EWAS catalog, we can identify proteins whose levels correlate with age and those that anti-correlate with age.

The first group includes proteins whose levels anti-correlate with age-protective CpGs, and proteins that correlate with age-damaging CpGs. For example, age-damaging cg19588519 located within the CHST15 gene is associated with higher levels of C-reactive protein, an inflammatory biomarker that is an important risk factor associated with age and aging-related diseases. Another example of an age-correlated protein is ECM1 whose levels are negatively related to higher methylation of an age-protective cg15654264 CpG site within the RPRD2 gene. ECM1 contributes to the maintenance of skin integrity and homeostasis. Its expression in human skin depends on age and UV exposure. Intrinsically (UV-protected) aged skin shows a significantly reduced expression of ECM1 in basal and upper epidermal cell layers compared with young skin. However, in photoaged skin, ECM1 expression is significantly increased within the lower and upper epidermis compared with age-matched UV-protected sites.

Proteins that anti-correlate with age contain proteins that correlate with age-protective CpGs and those that anti-correlate with age-damaging proteins. For example, age-protective cg21627412 is associated with lower levels of the collagen protein COL2A1 which has been reported to be markedly reduced in the aged dermis. This age-protective CpG site also protects from melanin hyperpigmentation (*b* = − 7.4, *p* = 0.016) and other disorders of pigmentation (*b* = − 5.84, *p* = 0.007). Another example is an age-damaging CpG (cg14036479) that correlates with levels of Biotinidase (BTD) which is involved in biotin metabolism.

Connecting CpGs with proteins results in a sparse CpG-protein-facial aging network that contains 90 damaging proteins and 61 protective proteins. Overlapping proteins, removed from further analysis, are likely due to false positives in the results of EWMR, or EWAS. We performed the functional analysis on the level of genes and proteins using the FUMA tool which yielded several insights into skin aging-related categories [Table S3].

### 3.4 Causal CpGs and skin pigmentation

Several top CpGs causal to facial aging are located within genes that have roles in skin pigmentation. Specifically, there are 6 protective CpGs in the IRF4 gene whose (hyper)methylation is highly significantly associated with lower facial age. The interferon regulatory factor, IRF4, is expressed in melanocytes in the skin. It is strongly associated with human pigmentation, sensitivity of skin to sun exposure, freckles, blue eyes, and hair color. The EW-GE MR analysis identifies an age-protective CpG (cg26422851) in the IRF4 gene that may causally lower expression of this gene (*b* = − 9.4, *p* = 4 *×* 10 − 5 that in turn, as found by GEMR, may have a protective effect on facial aging (*b* = 0.2, *p <* 10^−8^). Hence, this age-protective CpGs in the IRF4 gene is likely to slow facial aging via lowering expression of the IRF4 gene.

One of the top damaging methylation markers (cg14044465) is found in the BNC2 gene, while another CpG (cg03291755) in this gene is an anti-aging candidate marker that also causally lowers its expression (*b* = − 11.8, *p* = 4.8 *×* 10^−8^). The BNC2 gene has been reported to be associated with skin color variation in Europeans [28] and East Asians [29]. Interestingly, our EWMR analyses further identify that the damaging CpG site in the BNC2 gene (cg14044465) causally increases the odds of actinic keratosis (*b* = 0.36, *p* = 0.0003), while the age-protective CpG cg03291755 has a protective effect against actinic keratosis (*b* = −3.64, *p* = 4.5 *×* 10^−6^), and basal cell carcinoma (*b* = −2.12, *p* = 3.72 *×* 10^−7^).

Furthermore, both IRF4 and BNC2, as well as RALY/ASIP are implicated in facial pigmented spots during aging through pathways independent of basal melanin production [30]. The ASIP gene harbors two damaging causal CpGs and an age-protective causal CpG (cg05267394) that may also causally protect from facial pigmentation (*b* = − 19.65, *p* = 0.048), basal cell carcinoma (*b* = − 3.49, *p* = 0.003), and actinic keratosis (*b* = − 7.67, *p* = 0.0002).

Another pigmentation-related gene worth mentioning is the EXOC2 gene which is associated with hair color, skin pigmentation, and other pigmentation phenotypes [31]. Our EWMR analysis finds multiple protective and damaging CpGs in the EXOC2 gene. Specifically, two protective CpGs (cg20223322, cg26187313) are also identified to causally lower the odds of basal carcinoma (*b* = − 3.58, *p* = 0.0008), while a damaging methylation marker (cg17645074) in the EXOC2 gene increases the odds of this skin cancer (*b* = 2.26, *p* = 0.0013). While the MC1R gene coding for melanocortin 1 receptor has been dubbed a ‘youthfulness’ gene [32], and found to contribute to facial pigmented spots as well [30], our analysis did not identify causal CpGs in this gene.

Overall, the EWMR analysis of multiple skin aging-related phenotypes confirms that age-protective CpGs in skin pigmentation genes generally protect against age-related skin conditions and skin cancer, while age-damaging CpGs contribute to the risks. Functional analysis, using the FUMA tool, of all annotated genes associated with causal CpGs revealed chr16q24 as highly significantly (*p* = 3.4 *×* 10^−^9) over-represented positional set that contains 17 genes [Table S3]. Remarkably, the region chr16q24.3 was identified to be associated with skin pigmentation and skin aging in a recent comprehensive meta-analysis of 44 GWAS and gene expression studies [33]. This region is also over-represented in genes associated with protective genes (*p* = 0.0002) [Table S3].

Further, the facial pigmentation GWAS trait is also over-represented (*p* = 0.004) in all genes with causal CpGs. Other pigmentation-related over-represented traits include tanning (*p* = 0.012 based on three genes ASIP, IRF4, EXOC2) and low tan response (*p* = 10^−5^ based on ASIP, BNC2, IRF4, SPIRE2, AFG3L1P, DBNDD1, TRPS1). Further, causal genes are enriched in skin cancer-related traits, such as basal cell carcinoma (*p* = 3.4 *×* 10^−5^), cutaneous squamous cell carcinoma (*p* = 7.6 *×* 10^−5^), non-melanoma skin cancer (*p* = 0.01), and melanoma (*p* = 0.016).

### 3.5 Causal CpGs and extracellular matrix

The most visible hallmark of aging skin is the formation of wrinkles due to the loss of skin elasticity. Our unbiased EWMR analysis identified several causal CpG sites in genes with functions in elastic fiber formation, collagen biosynthesis and modifying enzymes, ECM glycoproteins, and semaphorin interactions.

Specifically, a highly protective CpG (cg05010648) in the ELN gene coding for the elastin protein has a large protective effect on facial aging (OR= 0.59). While currently there is no information on the downstream effect of this CpG on the level of elastin, it is a good candidate for an age-protective methylation marker.

A number of causal CpGs in two enzymes, LOX and LOXL, responsible for elastin cross-linking are also identified. While the elastin mRNA level generally remains quite stable at all ages, the levels of lysyl oxidase LOX and lysyl oxidase-like LOXL are affected by age and therefore have been suggested as targets to induce elastogenesis in adult skin [34]. There are 9 age-protective CpGs within the LOX gene, and the GEMR analysis identified that higher expression of the LOX gene in both exposed and not-exposed skin is causally associated with slightly lower facial age. Another related result is that a damaging CpG (cg02812767) causally increases expression of the LOXL1 gene (*b* = 6.47, *p* = 0.0009).

The EWMR analysis also identifies an age-protective methylation marker (cg08464402) in the collagen gene COL8A1. Extending annotations to the protein space, via the EWAS catalog, points to an additional age-protective CpG (cg21627412) that increases levels of the collagen protein (COL2A1), while a damaging CpG (cg13280184) correlates with levels of the MMP3 enzyme which is known to get upregulated with age and sun exposure. The MMP3 enzyme degrades ECM proteins, such as type IV, V, IX, and X collagens, gelatin, fibrillin-1, fibronectin, laminin, and proteoglycans, contributing to photoaging, wrinkle formation, and skin laxity.

These are potentially important findings on causal methylation markers that may slow down skin aging via higher levels of collagen proteins (e.g. COL8A1) or accelerate aging via reducing levels of collagen proteins (e.g. COL2A1) or increasing rate of collagen degradation (e.g. MMP3).

Functional analysis revealed significantly over-represented ECM-related gene-sets in the list of genes with causal CpGs [Table S3]. These gene-sets include ECM organization (*p* = 0.0006 based on 15 genes, including COL8A1, DAG1, DDR1, EFEMP1, ELN, FN1, GDF5, LAMB2, LOX, LOXL1, LTBP2, MMP2, P4HA2, SDC1, TNXB); elastic fibre formation (*p* = 0.0036); ECM glycoproteins (*p* = 0.0044; BGLAP, CRELD2, EFEMP1, ELN, FN1, LAMB2, LTBP2, POMZP3, PXDNL, RSPO2, SLIT1, THBS2, TNXB, VWA7). Further, causal genes are significantly enriched in semaphorin receptor binding (*p* = 0.04; SEMA6C, SEMA4B, SEMA4C, SEMA4D) and response to wounding (*p* = 0.0004; based on 31 genes) [Table S3].

Similar results are obtained on the protein level. For example, age-correlated proteins contain 16 proteins with general matrisome functions (NABA MATRISOME, *p* = 0.01; ECM1, F13B, LEFTY2, HPX, MMP3, IGF1, SERPINA12, SERPINF2, CCL11, REG3G, CST8, CXCL11, ESM1, SFTA2, PLXNA4, SVEP1), while anti-correlated proteins are enriched in collagen containing ECM (*p* = 0.009; PRG2, COL2A1, POSTN, SERPINA1, GH1, SERPINE2, PTN, L1CAM) as well as wound healing (*p* = 0.011; MPL, CD44, POSTN, SERPINA1, NOG, SERPINE2, LARGE, HMOX1, KLKB1) [Table S3].

### 3.6 Causal CpGs and Other Skin Aging Processes

There are multiple causal CpGs in other genes or proteins with known roles in skin health and skin aging. Below, we consider two skin-related processes, fatty acid metabolism, and oxidative stress.

Firstly, there are multiple causal CpGs in the FADS1 and FADS2 genes that code for fatty acid desaturases (FADS) with critical roles in fatty acid metabolism. Fatty acids impact a wide range of biological activities, including immune signaling and inflammation. They contribute 15% of the total lipid content within the skin surface lipids that regulate skin barrier permeability and modulate the risk of infection by interacting with skin microbiota. FADS have been reported to impair skin barrier function contributing to skin pathology and reducing total lipid content in the skin of older adults. This is believed to contribute to dryness, increased susceptibility to infection, and impaired wound healing in response to injury. A recent study demonstrated that the phenotypic associations of rs174537 are likely due to methylation differences that may directly impact FADS activity [35]. Specifically, rs174537 is anti-correlated with cg27386326 which is identified as an age-protective causal CpG in the EWMR analysis.

Another study [36] examined the relationship between keratinocyte fatty acid composition, lipid biosynthetic gene expression, gene promoter methylation, and age. They found that expressions of FADS genes and elongases (ELOVL6 and ELOVL7) were lower in adult versus neonatal keratinocytes, and were associated with lower concentrations of n-7, n-9, and n-10 polyunsaturated fatty acids in adult cells. Hence, causal CpGs in the FADS cluster represent potentially actionable methylation markers for treating early signs of skin aging. A recent MR analysis reported causal associations between genetically proxied unsaturation degree of fatty acids and the pace of facial skin aging [37].

Several GWAS traits related to fatty acids are significantly over-represented in genes with CpGs causal to facial aging, including plasma omega-3 polyunsaturated fatty-acid levels (docosapentaenoic acid, eicosapentaenoic acid, alpha-linolenic acid); plasma omega-6 polyunsaturated fatty-acid levels (adrenic acid, arachidonic acid) and very long-chain saturated fatty-acid levels (fatty acid 22:0) [Table S3].

Another skin aging-related category is increased oxidative stress. The oxidative stress theory of aging states that age-associated functional losses are due to the accumulation of Reactive Oxygen Species (ROS). Overall, the endogenous antioxidant capacity (enzymatic and non-enzymatic) of the skin is lowered with age, and the aged skin is more vulnerable to external factors, especially UV radiation, pollution, and microorganisms. Since the epidermis is more exposed to external stimuli than the dermis, the ROS load is higher in the epidermis compared to the dermis [38].

One of the supporters of the antioxidant capabilities of the cell is glutathione. Age-related decline in the activity of the antioxidant enzyme glutathione peroxidase (GPX1) may contribute to increased free radicals and GPX1 was proposed as a potential therapeutic target for the enhancement of the regenerative capacity in old subjects [39]. The EWMR analysis identifies three causally age-protective CpGs (cg24011261, cg05551922, cg05055782) in the GPX1 gene. These CpGs are also found to be candidates for causal positive effects on the telomere length [Table S4] while at the same time increasing the odds of basal cell carcinoma (e.g. *b* = 4, *p* = 0.01 for cg24011261). Another xenobiotic enzyme that may lose its antioxidant capacity with age is NAT2. It is involved in the metabolism of drugs, environmental toxins, and carcinogens, and also affects levels of cholesterol and triglycerides. According to the EWMR analysis, NAT2 harbors two age-damaging CpGs that may be related to the decreased antioxidant capacity of aging skin.

Furthermore, genes with age-protective CpGs are significantly enriched in the cellular response to oxidative stress (*p* = 0.04; based on genes GPX1, ERO1L, MGMT, TPM1, PDE8A, MMP2, CBX8, NR4A2, PLA2R1, PARK2). Additionally, all causal genes are significantly enriched in the oxidation-reduction process (*p* = 0.0036; based on 36 genes) and the response to oxidative stress gene-set (*p* = 0.0096; based on 20 genes) [Table S3].

Among serum proteins correlated with lower perceived facial age, at least three aproteins (AKR1C3, ASL, HMOX1) have functions in xenobiotic metabolism. Overall, proteins correlated with more youthfulness are enriched in adhesion, response to bacteria, viruses, nutrients, growth factors, and acute inflammatory and immune response, while proteins correlated with older age are enriched in adaptive immune response, adaptive response to stress, as well as several metabolic pathways, including vitamin, phosphorus, carbohydrate, and small molecule metabolic processes [Table S3].

### 3.7 Causal CpGs and telomeres

Cellular senescence and telomere shortening are molecular hallmarks for aging in general and play important roles in skin aging. As senescent cells accumulate with age in various tissues, including skin, they secrete factors called the senescent-associated secretory phenotype (SASP) that trigger inflammation and remodel the extracellular matrix (ECM) contributing to premature aging phenotypes. Cell senescence is often approximated by a reverse of telomere length (TL) as shorter telomere length (more senescence) has been reportedly associated with age-related diseases and also with mortality.

However, the relationship between skin aging and TL is more complex. Specifically, longer telomeres have been reported to be associated with an increased risk of melanoma [40]. This link was first discovered because of a strong association between longer telomeres and a high number of naevi. In contrast, for cutaneous squamous cell carcinomas, the relationship is reversed with a higher risk in individuals with shorter telomeres.

We set out to explore the relationship between methylation markers causal to facial aging and TL. To this end, we ran the EWMR analysis on meQTLs from blood as exposure and TL as the outcome (see Methods). Firstly, there is a large overlap between CpGs causal to facial aging and CpGs causal to telomere length TL (235 CpGs). In contrast, the overlap between CpGs causal to facial aging and CpGs from the epigenetic estimator of telomere length (DNAmTL from the `methylCIPHER R` package) is very small (only 4 CpGs). It is worth noting that While the DNAmTL estimator has a high correlation with cellular replication rate, it contains only one CpG (cg00029246) in the DGKI gene that causally affects TL. This again points out that CpG sites in the DNAmTL estimator correlate with the length of telomeres and may be the consequences of telomere shortening rather than drivers of it.

Among common CpGs that are causal to both facial aging and TL, there are 19 age-damaging and TL-shortening CpGs located in the TNXB gene that encodes a member of the tenascin family of extracellular matrix glycoproteins with anti-adhesive effects. The TNXB gene plays a role in ECM organization and collagen fibrin formation. It also functions in matrix maturation during wound healing. Deficiency of TNXB has been associated with the connective tissue disorder Ehlers-Danlos syndrome. Six age-damaging CpGs in the TNXB gene are also associated with lower levels of the CRISP2 protein which is involved in cell-cell adhesion, regulation of ion channels’ activity, and is implicated in male fertility. Other common genes are GPSM3 and GPX1 with roles in regulating inflammatory response and protecting cells against oxidative damage. Specifically, the GPX1 gene that harbors three age-protective CpGs has a function in glutathione conjugation and fatty acid metabolism. Another gene in common is PSORS1C1 known to confer susceptibility to psoriasis and systemic sclerosis. It harbors an age-damaging and TL-lengthening CpG (cg01016122).

To further investigate complex relationships between epigenetic markers causal to facial aging and TL, we relaxed the p-value cut-off in the EWMR (*p* = 0.01) and obtained 486 CpGs causal to both traits [Table S4]. Interestingly, two groups of causal CpGs emerge in this dataset. The first group contains CpGs that exhibit behavior consistent with the prevailing views that short telomeres correlate with aging. This “consistent”group contains CpGs that may causally shorten TL and accelerate facial aging, or causally lengthen TL and slow down facial aging. The “reverse”group contains CpGs that causally affect TL and aging in opposite directions, e.g. longer TL and accelerated facial aging. Interestingly, correlations of the effect sizes from the EWMR analyses for facial aging and TL for the consistent and reverse groups taken separately are very strong (Pearson correlation coefficient in the consistent group = − 0.89, and in the reverse group = 0.87).

CpGs in the consistent group are significantly enriched in several processes related to skin development and maintenance including keratinization, keratinocyte differentiation, regulation of keratinocyte proliferation, epithelial cell differentiation, epidermal cell differentiation, cornified envelope, desmosome; vitamin D signaling pathway; responses to extracellular stimulus and nutrients, vitamins, interferon-gamma, and EGF [Table S4]. CpGs in the reverse group are significantly enriched in the regulation of signaling receptor activity, protein localization to cell surface, several metabolic processes (citrulline, arginine, nitric oxide, amino acids, glutamine), and pathways related to synaptic plasticity and transmission.

## 4 Discussion

The gigantic antiaging cosmetic industry has largely been focused on figuring out the most effective products that conceal, or temporarily mitigate the visible downstream effects of aging, rather than proactively address the upstream root causes of aging [41]. It has already been proposed that the prerequisite for an anti-aging rejuvenation treatment is a robust reduction in biological age, which can be assessed by biomarkers of aging, such as epigenetic clocks [42]. Skin-specific biological clock based on methylation markers [8] has already been used to identify a novel senotherapeutic peptide that improves skin health, reduces skin biological age, and decreases the expression of senescence markers [9].

However, the current generation of biological clocks is based on methylation sites that are correlated with chronological age, and hence their causal effect on skin aging is difficult to infer. To understand the causes of skin aging, and develop efficient senotherapeutic treatments, it is important to distinguish epigenetic drivers of aging from the consequences of aging. Furthermore, it is crucial to identify age-damaging methylation sites from age-protective ones.

In this study, we systematically evaluated the relationship between DNA methylation and facial aging. We utilized the EWMR procedure and identified hundreds of potentially causal methylation markers that affect facial aging. Subsequent functional analysis revealed several interesting insights. Firstly, it appears that age-protective, or adaptive, methylation changes are nearly as common as those methylation changes that drive, or accelerate, aging. This is consistent with a recent report on general aging [11]. Hence, identifying CpGs that causally impact youthfulness is as important as finding age-damaging causal CpGs. The development of rejuvenating skin treatments should target both groups of methylation markers to reverse damaging methylation sites on one hand while inducing, or maintaining, the protective methylation sites.

Secondly, our study reports little overlap between existing epigenetics-based biological clocks and CpGs causal to facial skin aging. One explanation is that causal damaging or adaptive CpGs may not change as much with age, and thus have not been included as input features in machine learning models for biological clocks. Indeed, consistent with an earlier study on general aging [11], we found no correlation between the size of the causal effect on facial aging and the magnitude of age-related changes.

Thirdly, our study demonstrates that methylation data from blood can be utilized to uncover damaging and protective CpGs causal to facial aging. Furthermore, identified causal CpGs shed light on biological processes that underlie facial skin aging and other dermatological traits. For example, age-protective CpGs in skin pigmentation genes generally protect against age-related skin conditions and skin cancer, while age-damaging CpGs contribute to the risks. Significant overlap in causal CpGs that contribute to facial aging and senescence further confirms that CpGs causal to aging underlie biological mechanisms that causally affect the pace of aging rather than those that are mere consequences of aging. At the same time, our study points out that the relationship between telomere length and facial aging is not linear, as some CpGs that contribute to longer telomeres may exert damaging effects on the pace of facial aging.

The limitation of this study is that we used a high p-value threshold in the EWMR procedure resulting in a higher proportion of false positives. The reason for this is to showcase the integration of blood methylation data with the summary statistics from GWAS on perceived facial age that is based on replies to the question of whether a person thinks others perceive her/him as looking younger or older than their real chronological age. It is worth noting that less than 5% of the UKBB participants reported “looking older than actual age”. Another limitation is that the perceived facial data is self-reported. Future studies should be based on more precise estimates of perceived facial age estimated by specialists such as dermatologists or aestheticians. Finally, this study utilizes the lumped data from females and males. Furthermore, it largely contains data from white British participants. To understand the epigenetics of facial aging future studies on should be based on separate analyses of female and male datasets as well as consideration of other ethnicities.

Overall, this is the first study that uncovers epigenetics markers causal to facial aging. It provides proof-of-concept for utilizing blood-measured CpGs in elucidating biological processes that underlie facial aging. Novel causal CpGs can be utilized as input features for building the biological clock of facial aging, predicting personalized face aging trajectories (ageotypes), and developing novel rejuvenating topical, and ‘beauty from within’ senotherapeutic solutions that target root causes and early signs of aging.

